# Molecular Docking studies of Phytocompounds of *Rheum emodi* Wall with proteins responsible for antibiotic resistance in bacterial and fungal pathogens: *In silico* approach to enhance the bio-availability of antibiotics

**DOI:** 10.1101/2020.05.10.086835

**Authors:** Rajan Rolta, Vikas Kumar, Anuradha Sourirajan, Kamal Dev

## Abstract

*Rheum emodi* Wall. (Himalayan rhubarb) has been used to cure many human diseases. Literature survey demonstrated that it has many pharmacological activities such as antioxidant, antimicrobial, antiviral, anticancer and wound healing. The present study was aimed to understand if major phytocompounds of *Rheum emodi* could bind proteins responsible for antibiotic resistance in bacterial and fungal pathogens and enhance the potency of antibiotics. The major phytocompounds of *R. emodi* (emodin, rhein-13c6 and chrysophenodimethy ether) were retrieved from Pubchem and target proteins were retrieved from RCSB protein data bank. The docking study was performed with Hex 8.0.0 software and molinspiration, swiss ADME servers were used for determination of Lipinski rule of 5, drug-likeness prediction respectively, whereas, admetSAR and Protox-II tools were used for toxicity prediction. Among all the selected phytocompounds, emodin showed the best binding energy of −235.82 Kcal mol^-1^ and −245 Kcal mol^-1^ with cytochrome P450 14 alpha-sterol demethylase (PDB ID: 1EA1) and **N-**myristoyl transferase (PDB ID: 1IYL) receptors, respectively, which is more than that of fluconazole (−224.12 kcalmol^-1^ and −161.14 kcal mol^-1^). Similarly, with Penicillin binding protein 3 (PDB ID: 3VSL) receptor, emodin and Chrysophanol dimethyl ether showed highest binding energy of - 216.68 Kcal mol^-1^ and −215.58 kcal mol^-1^ which is comparable to erythromycin (−263.63 kcal mol^-1^), chloramphanicol (−217.34 kcal mol^-1^) and tetracycline (−263.63 kcal mol^-1^). All the selected phytocompounds also fulfill Lipinski rule, non-carcinogenic and non-cytotoxic in nature. These compounds also showed high LD_50_ value showing non-toxicity of these phytocompounds.

**Graphical abstract:** 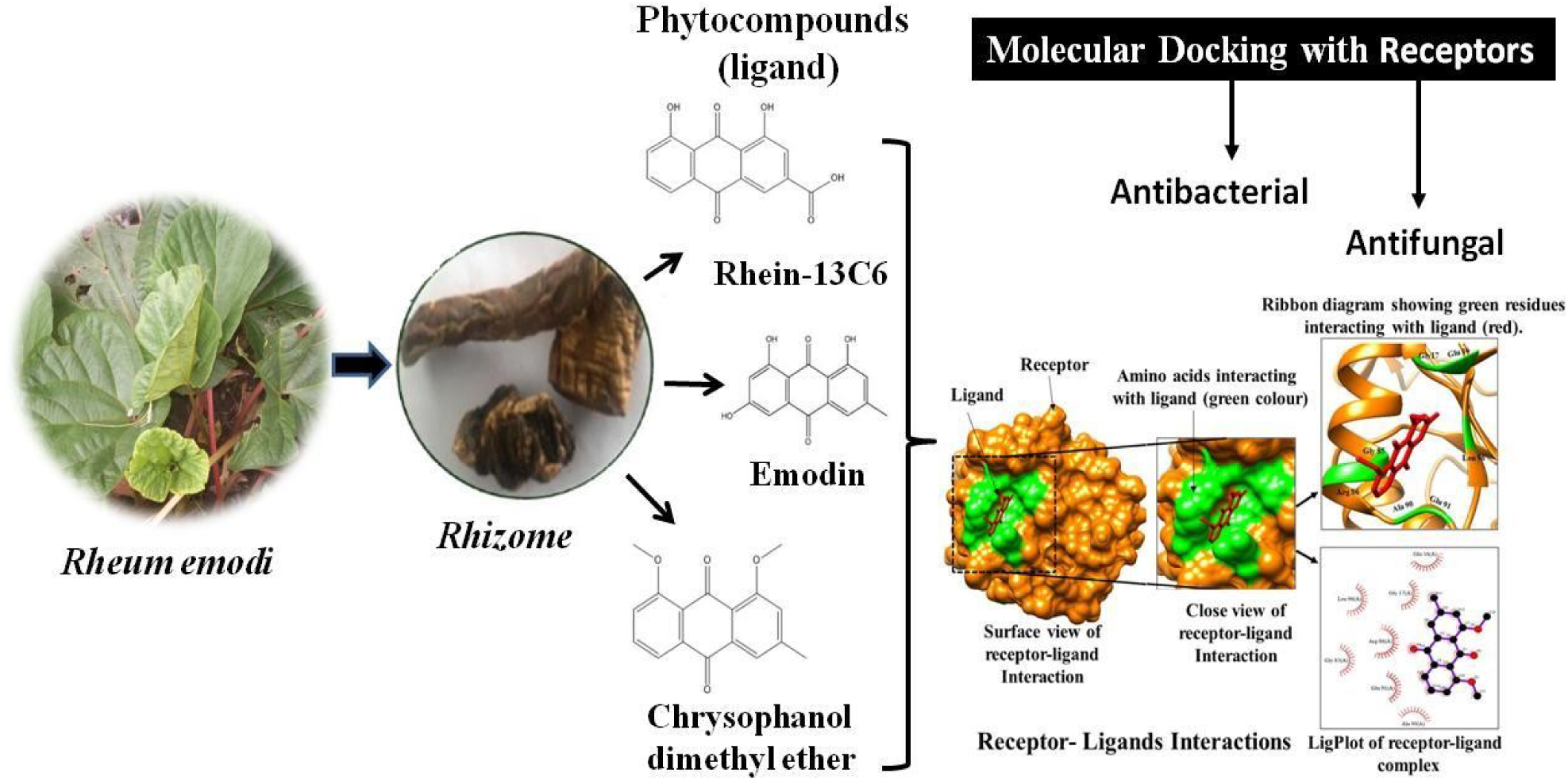

## 1. Introduction

Since earliest times, many plants have been known to exert healing properties against human infections due to their content of secondary metabolites, which in more recent times have been found to act as antimicrobial agents against human pathogens. Over the past decade, much attention has been placed on the study of phytochemicals for their antibacterial activity, especially against multidrug-resistant Gram-negative and Gram-positive bacteria (Borges *et al.*, 2015). Now a days, antimicrobial resistance is a major global problem caused by bacterial and fungal strains. In the past two decades, acquired MDR infections have increased due to the production of β-lactamases (e.g. extended spectrum β-lactamases [ESBLs] enzymes, carbapenemases, and metallo-β-lactamases), leading to third generation cephalosporin and carbapenem resistance (Blair *et al.*, 2015). Mechanism of drug resistance is classified in three categories modification in enzyme, mutation in antibiotics and increase in the activeness of efflux pump. Efflux pump transporter are present in all organisms eukaryotes and prokaryotes, it extrude a verity of compounds and chemicals from the cells (Zgurskaya and Nikaido, 2000; Ramos *et al.*, 2002). McMurry *et al.* (1980), first time reported that bacteria can acquire the antibiotic resistance by extruding the antibiotics. Efflux pump are grouped in the five structural families namely the resistance nodulation-division (RND) (Tseng *et al.*, 1999), the small multidrug resistance (SMR) (Chung *et al.*, 2001), the multi antimicrobial extrusion (MATE) (Kuroda *et al.*, 2009), the major facilitator superfamily (MFS) (Law *et al.*, 2008), and the ATP-binding cassette (ABC) (Lumbelski *et al.*, 2007) superfamilies. Drug resistance pathogens will infect more than 444 million people on the globe by the year 2050 (Bartlett *et al.*, 2013; Gould and Bal, 2013; Aslam *et al.*, 2018). The results emergence of multidrug resistance bacterial and fungal strains needs rapid development of new antimicrobial drugs to combat drug resistance. Plant-based phytochemicals offer attractive, effective, and holistic drug action against the pathogens without much of the side effects.

Molecular docking plays an important role in the rational design of drugs. In the field of molecular modeling, docking is a method to predicts the preferred orientation of one molecule to a second when bound to each other to form a stable complex. Molecular docking can be defined as an optimization problem, which would describe the “best-fit” orientation of a ligand that binds to a particular protein of interest (Lengauer and Rarey, 1996; Kumar *et al*., 2018). Medicinal plants play important role to cure different types of diseases. Traditional Medicine (TM) may offer an abundance prospect to combat drug resistance (Gupta and Birdi, 2017; Ahmad and Beg, 2001). *Rheum emodi* Wall. is one of the important medicinal herbs of Chinese medicinal system (Singh *et al.*, 2017). Rolta *et al.* (2018) reported the antioxidant, antibacterial, antifungal activities of methanolic extracted of *R. emodi* rhizome. Anthraquinones including rhein, chrysophanol, aloe-emodin, emodin, physcion (emodinmonomethyl ether), chrysophanein and emodin glycoside are the major phytocompounds of *R. emodi* (Malik *et al.*, 2010; Malik *et al.*, 2016). Anthraquinones has various pharmacological properties including anti-inflammatory, antimicrobial, antimicrobial, antimutagenic, immunomodulatory, and synergistic activity (Malik and Muller, 2016; Sharma *et al.*, 2017: Rolta *et al.*, 2020). In our previous study, we reported that emodin, emodin D4, rhein-13C6, Resveratrol and chrysophanol dimethyl ether in chloroform sub fraction of methanolic extract of *R. emodi* rhizome emodin showed profound synergistic activity in combination with antibacterial and antifungal antibiotics and lowered the dosage of antibiotics by 4–257 folds (Rolta *et al.*, 2020). The mechanism of synergistic potential of phytocompounds is complex and still remined unanswered. The possible mechanisms are alternation of host targets responsible for drug resistance, modification of antibiotics such that they are no longer sensitive to effectors of drug resistance, blocking efflux of antibiotics etc. To understand the mechanisms of synergistic potential of phytocompounds of *R. emodi*, the present study was designed to study the *In-silico* binding of phytocompounds of *R. emodi* with bacterial and fungal proteins responsible for inactivation of bacterial and fungal antibiotics and lead to drug resistance.

## 2. Material and methods

### 2.1 Bioinformatics tools

Various bioinformatics tools used in the present study are Hex 8.0.0 software (http://hex.loria.fr/dist/index.php), Open Babel GUI (O’Boyle et al., 2011), Molispiration(https://www.molinspiration.com/),admetSAR(http://lmmd.ecust.edu.cn/admetsar1/predict), PROTOX-II (http://tox.charite.de/protox_II/), Chimera 1.8.1 (Pettersen *et al*., 2004) and LigPlot (Laskowski and Swindells, 2011).

### 2.2 Protein preparation

The 3D crystal structures of selected target proteins (Table-1) responsible for antibacterial and antifungal potential were retrieved from RCSB PDB (http://www.rscb.org/pdb). All proteins had co-crystallized ligands (X-ray ligands) in their binding site. These complexes bound to the receptor molecule, such as non-essential water molecules, including heteroatoms were removed from the target receptor molecule. Finally, hydrogen atoms were added to the target receptor molecule.

### 2.3 Ligand preparation

Three phytocompounds namely emodin, Chrysophanol dimethyl ether, and rhein-13C6 were selected based on results of our previous study (Rolta *et al.*, 2020) and are further investigated to study the molecular mechanism of their interactions with bacterial and fungal pathogens. Antibiotics such as tetracycline, chloramphanicol, erythromycin, fluconazole, amphotericin B were used as standard control. The 2-dimensional structures of all the phytocompounds and antibiotics were obtained from Pubchem (www.pubchem.com) in .sdf format. The .sdf file of phytocompounds was converted into PDB format by using Open Babel tool (Wang *et al*., 2009; Noel *et al.*, 2001). Molecular structures and weight of selected phytocompounds are listed in table-2.

**Table- 1:**
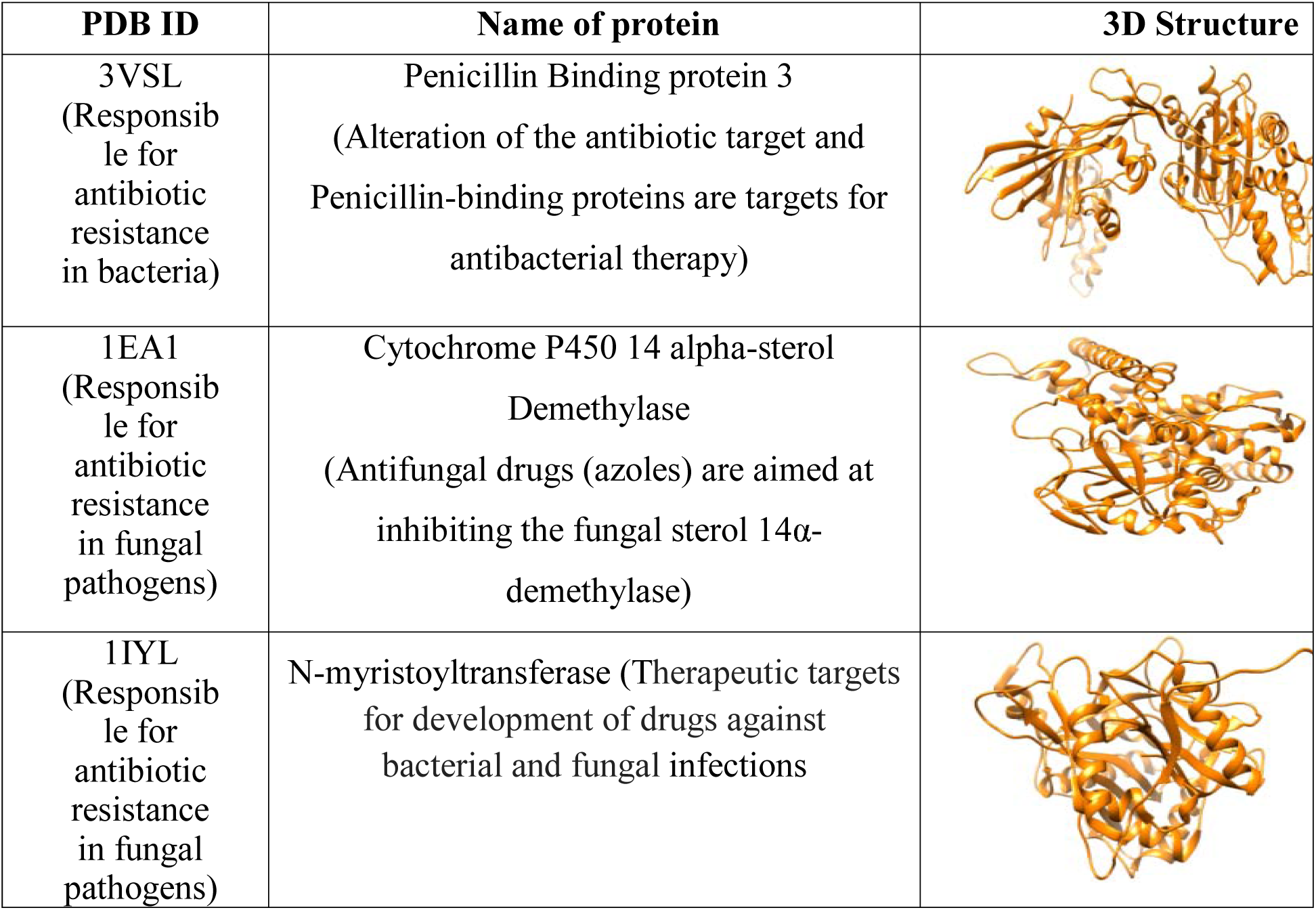
Target receptor proteins responsible for antibiotic resistance in bacteria and fungi.

| PDB ID | Name of protein | 3D Structure |
| --- | --- | --- |
| 3VSL<br>(Responsible for antibiotic resistance in bacteria) | Penicillin Binding protein 3<br>(Alteration of the antibiotic target and Penicillin-binding proteins are targets for antibacterial therapy) |  |
| 1EA1<br>(Responsible for antibiotic resistance in fungal pathogens) | Cytochrome P450 14 alpha-sterol Demethylase<br>(Antifungal drugs (azoles) are aimed at inhibiting the fungal sterol 14α-demethylase) | |
| 1IYL<br>(Responsible for antibiotic resistance in fungal pathogens) | N-myristoyltransferase (Therapeutic targets for development of drugs against bacterial and fungal infections) |  |

**Table- 2:**
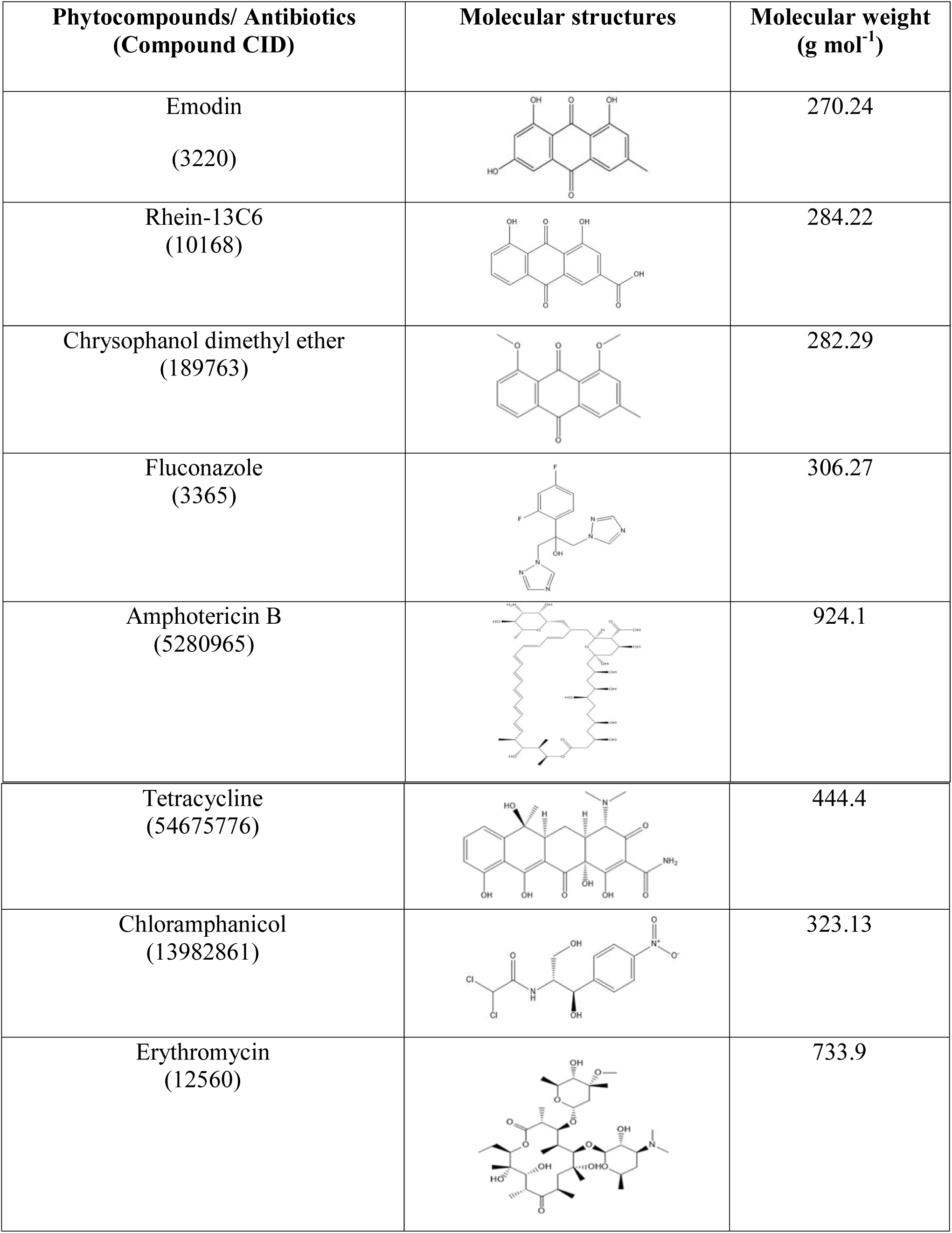
Molecular structure, molecular weight, and CID no. of selected phytocompounds and standard drugs.

| Phytocompounds/ Antibiotics<br>(Compound CID) | Molecular structures | Molecular weight<br>(g mol<sup>-1</sup>) |
| --- | --- | --- |
| Emodin<br>(3220) |  | 270.24 |
| Rhein-13C6<br>(10168) |  | 284.22 |
| Chrysophanol dimethyl ether<br>(189763) |  | 282.29 |
| Fluconazole<br>(3365) |  | 306.27 |
| Amphotericin B<br>(5280965) |  | 924.1 |
| Tetracycline<br>(54675776) |  | 444.4 |
| Chloramphenicol<br>(13982861) |  | 323.13 |
| Erythromycin<br>(12560) |  | 733.9 |

### 2.4 Docking of receptor with ligands

The docking of selected ligands to the catalytic pocket of protein was performed using Hex 8.0.0. The docking complex was generated after the completion of docking and saved as .pdb file. The .pdb complex of protein and phytocompounds were further analyzed by PDBsum (www.ebi.ac.uk/pdbsum) to study the list of interactions between target proteins and phytocompounds. Detailed visualization and comparison of the docked sites of target proteins and ligands were done by Chimera (Pettersen *et al*., 2004) and LigPlot (Laskowski and Swindells, 2011).

### 2.5 Drug likeness calculations

Drugs scans were carried out to determine whether selected phytochemicals fulfill the drug-likeness conditions. Lipinski’s filters using Molinspiration (http://www.molinspiration.com/) were applied for examining drug likeness attributes as including quantity of hydrogen acceptors (should not be more than 10), quantity of hydrogen donors (should not be more than 5), molecular weight (mass should be more than 500 daltons) and partition coefficient log P (should not be less than 5). The smiles format of each of the phytochemical was uploaded for the analysis (Rosell and Crino, 2002).

### 2.6 ADMET screening and toxicity prediction of phytocompounds and standard antibiotics

ADMET screening was done to determine the absorption, toxicity, and drug-likeness properties of selected phytocompounds. The 3D structures of phytocompounds (emodin, chrysophanol dimethyl ether, rhein-13C6) and standard drugs (chloramphenicol, erythromycin, tetracycline, fluconazole and amphotericin B) were saved in .smiles format and drug were uploaded on admetSAR (Laboratory of Molecular Modeling and Design, Shanghai, China), and PROTOX-II webservers (Charite University of Medicine, Institute for Physiology, Structural Bioinformatics Group, Berlin, Germany). The admet SAR provides ADMET profiles for query molecules and can predict about fifty ADMET properties. Toxicity classes are as follows: (i) Category I contains compounds with LD50 values ≤50 mg kg^-1^, (ii) Category II contains compounds with LD50 values >50 mg kg^-1^ but 500 mg kg^-1^ but 5000 mg kg^-1^ (Cheng *et al.*, 2012; Yang *et al.*, 2019). PROTOX is a Rodent oral toxicity server predicting LD50 value and toxicity class of query molecule. The toxicity classes are as follows: (i) Class 1: fatal if swallowed (LD50 ≤5), (ii) Class 2: fatal if swallowed (55000) (Banerjee *et al.*, 2018).

## 3. Results

### 3.1 Receptor-ligands interactions

The results of docking interactions between selected phytocompounds and targeted receptor proteins were shown in Fig. 1, 2 and Table 3. It was found that emodin showed best interaction with penicillin binding protein 3 (3VSL) with docking score (−216.68 kcal mol^-1^) followed by chrysophanol dimethyl ether (−215.58 kcal mol^-1^) which is comparative to chloramphanicol (−217.34 kcal mol^-1^) and lower than that of erythromycin (−263.63 kcal mol^-1^) and tetracycline (−263.63 kcal mol^-1^). Similarly, with N-myristoyl transferase, emodin showed highest binding energy (−245Kcal mol^-1^) followed by chrysophanol dimethyl ether (−230.88 kcal mol^-1^) as compared to that of fluconazole (−161.14 kcal mol^-1^). Chrysophanol dimethyl ether (−225.76 kcal mol^-1^) and emodin (−221 kcalmol^-1^) showed higher binding energy with cytochrome P450 14 alpha-sterol demethylase which is comparable with that of fluconazole (−224.12 kcalmol^-1^). Amphotericin B showed higher binding energy with both N-myristoyl transferase (−362.43 kcalmol^-1^) and cytochrome P450 14 alpha-sterol demethylase receptors (IEA1) (−387.92 kcalmol^-1^) (Table-3). The interacting amino acids showing H-bonding and hydrophobic interactions between phytocompounds and receptors are shown in table-3. Interactions of various amino acids of antibacterial receptor proteins (3VSL) and antifungal proteins (1EA1 and 1IYL) phytocompounds were visualized through chimera 1.8.1 and LigPlot analysis as shown in Fig. 1-3.

**Table 3:**
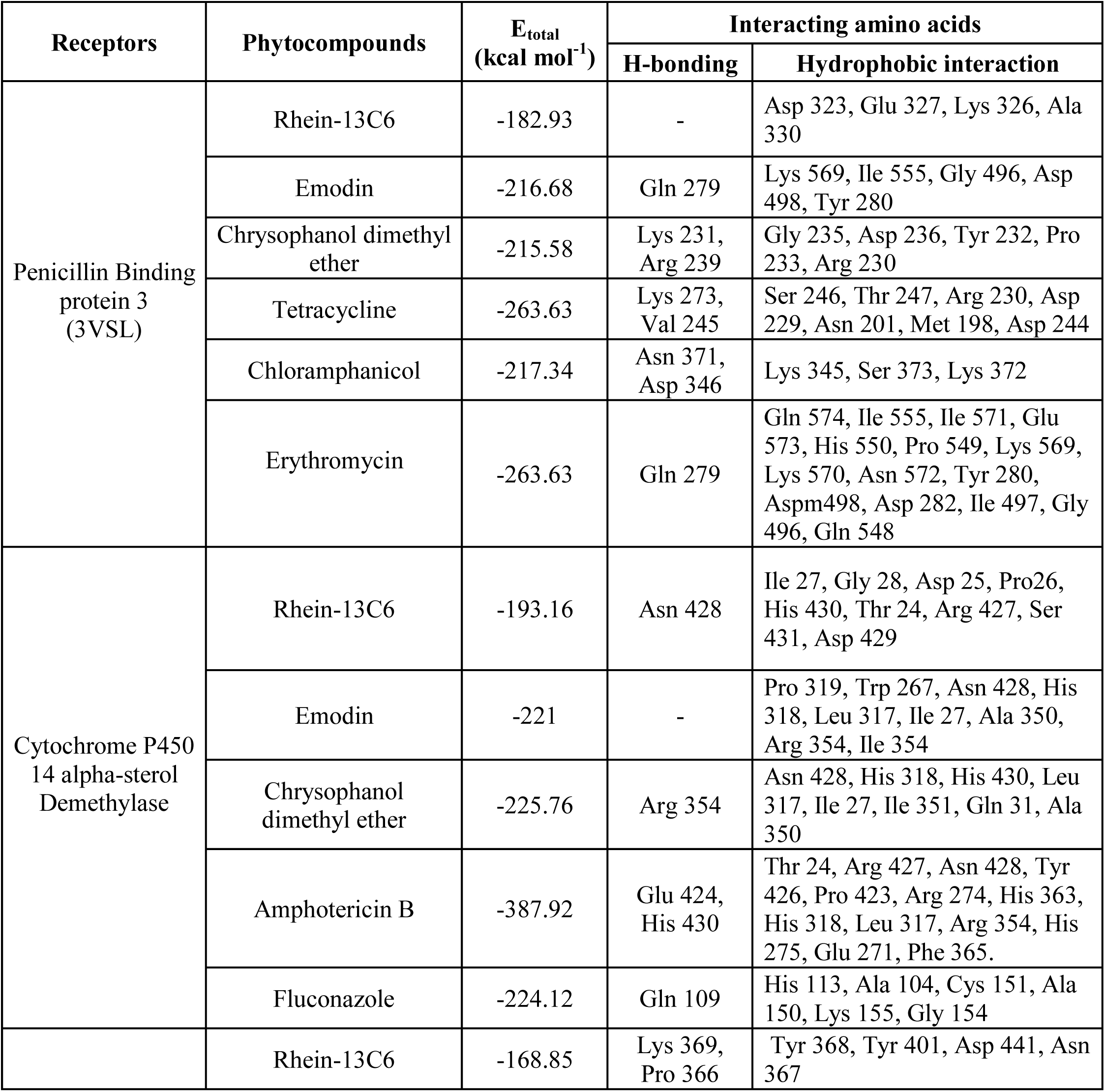
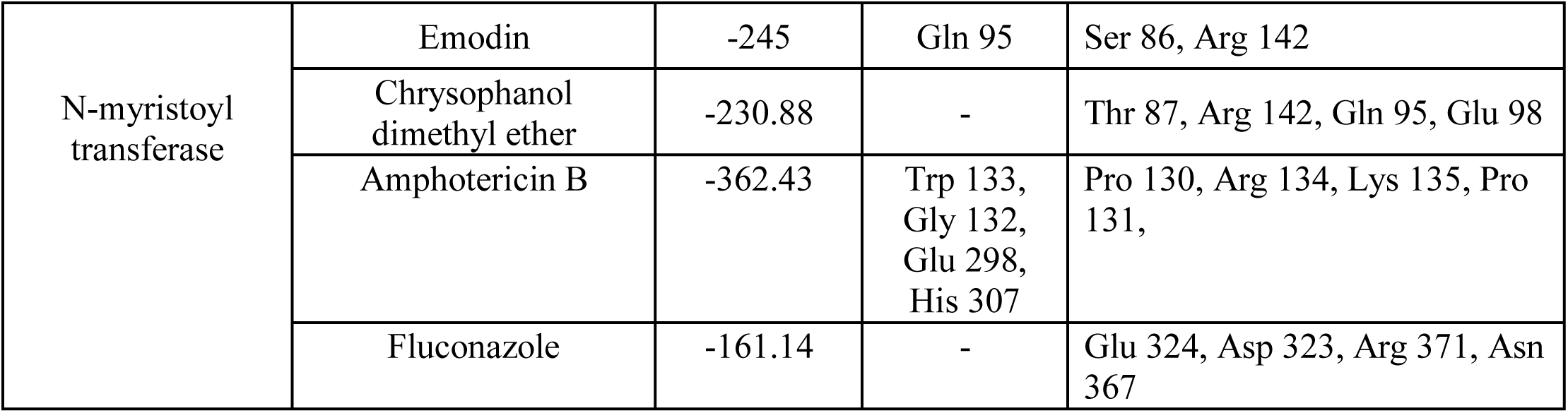
E-total of selected phytocompounds of *R. emodi* and antibiotics with bacterial and fungal targets.

**Fig. 1:**
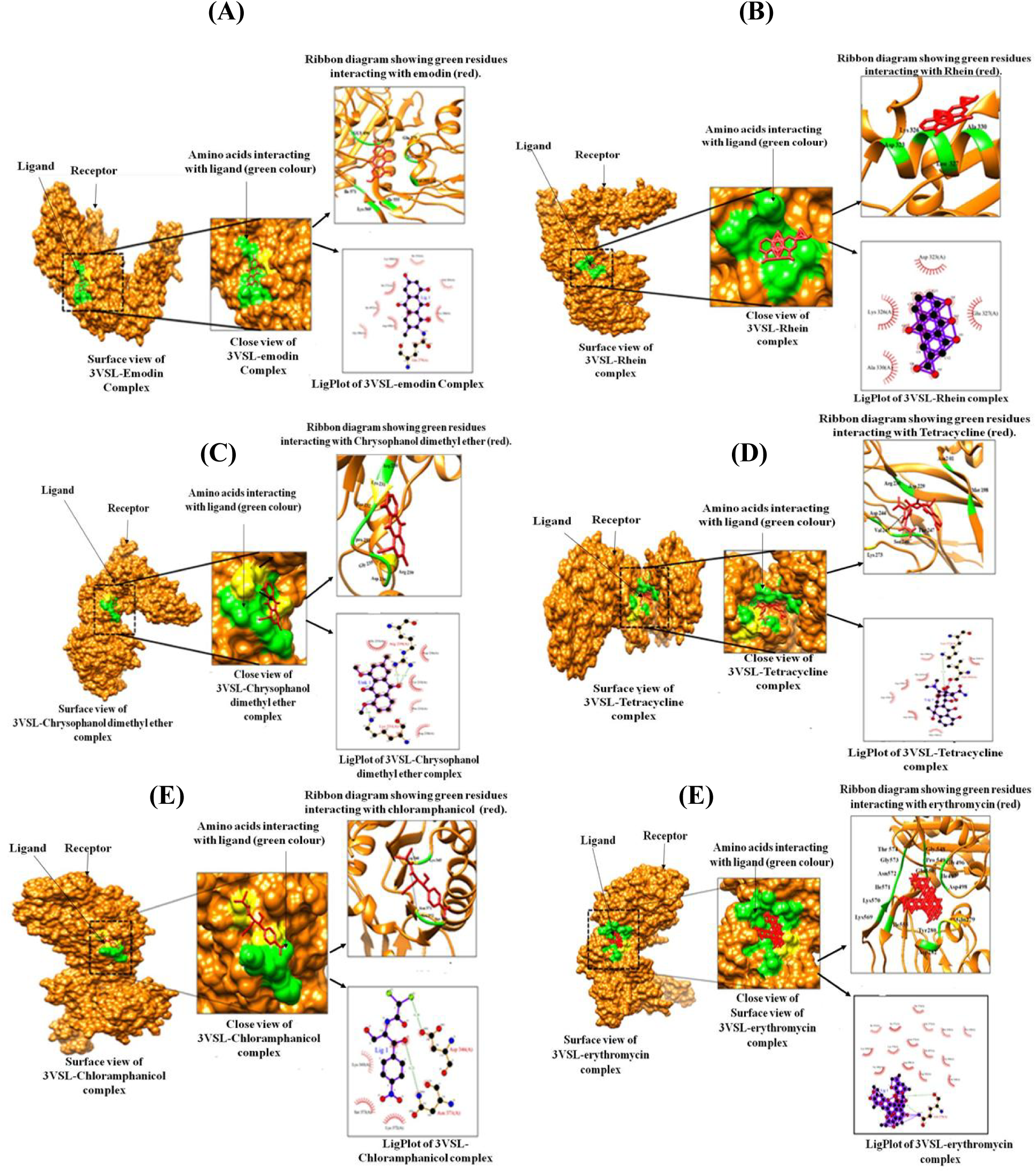
Interactions of phytocompounds with bacterial receptors Penicillin Binding protein 3 (3VSL). Interactions of emodin (A), Rehin-13C6 (B), Chrysophanol dimethyl ether (C), Tetracycline (D), Chloramphanicol (E), and erythromycin (F). Each panel shows surface view of receptor-ligand complex, close view of receptor-ligand complex, Ribbon diagram of receptor showing green residues interacting with ligand (red), and LigPlot of receptor-ligand complex as indicated.

**Fig. 2:**
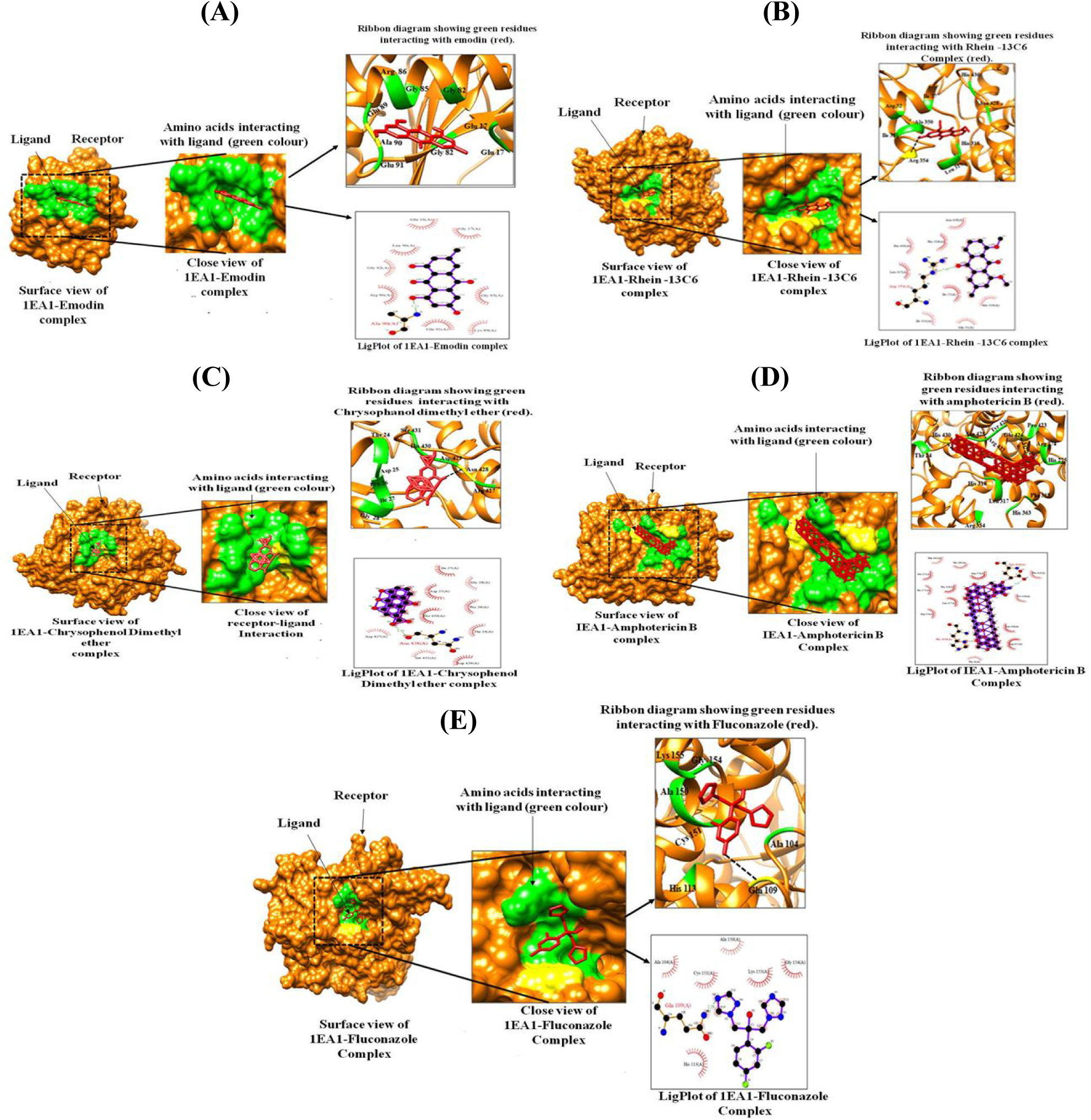
Showing interactions of phytocompounds with fungal receptors Cytochrome P450 14 alpha-sterol demethylase (PDB ID: 1EA1). Interactions with emodin (A), interactions with Rhein-13C6 (B), interactions with Chrysophanol dimethyl ether (C), interactions with Amphotericin B (D), and interaction with fluconazole (E). Each panel shows surface view of receptor-ligand complex, close view of receptor-ligand complex, Ribbon diagram of receptor showing green residues interacting with ligand (red), and LigPlot of receptor-ligand complex as indicated.

**Fig. 3:**
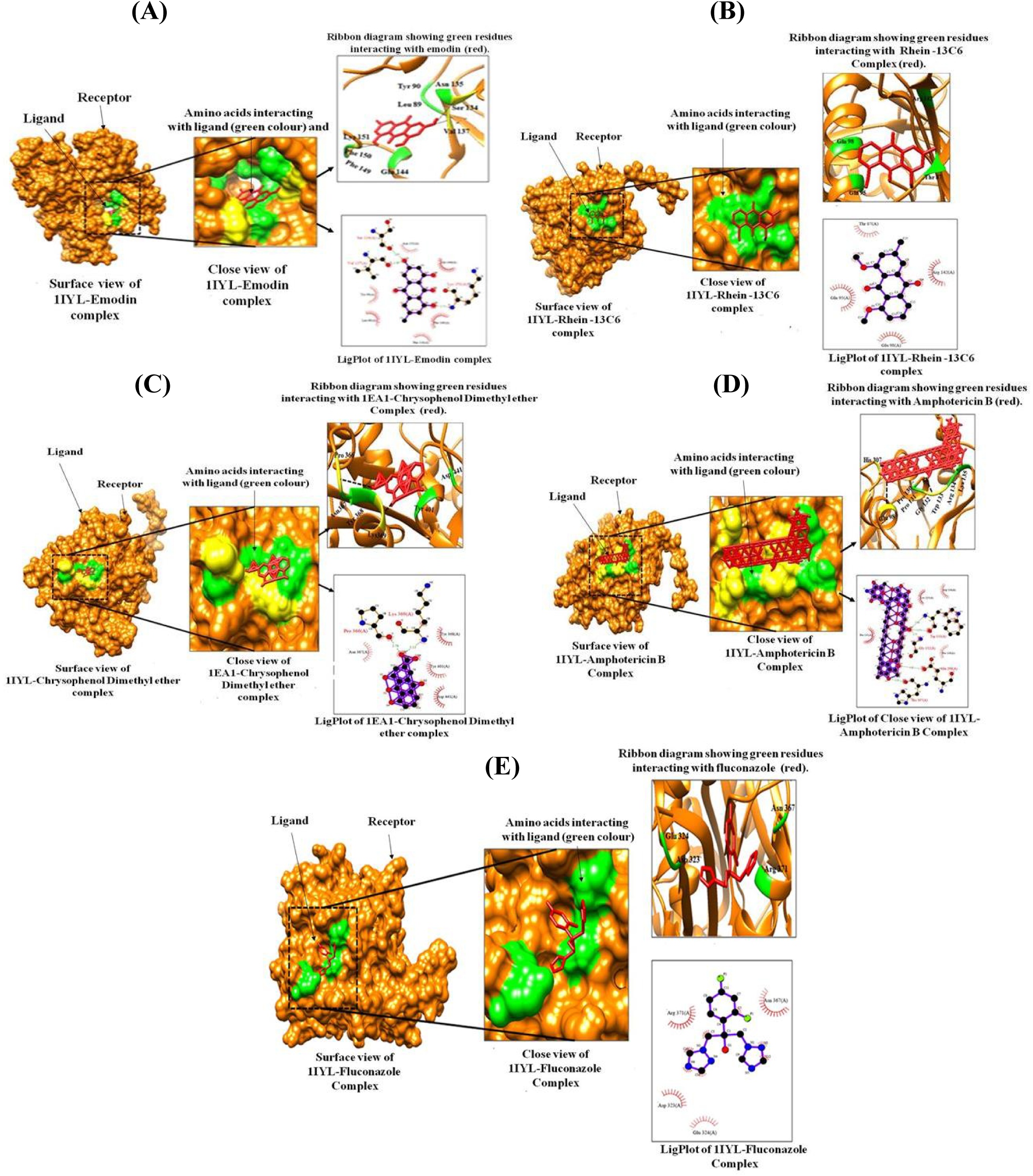
Showing interactions of phytocompounds with fungal receptor, NMT-Mysritel transferase (PDB ID: 1IYL): Interactions with emodin (A), interactions with Rhein-13C6 (B), interactions with Chrysophanol dimethyl ether (C), interactions with Amphotericin B (D) and interactions with fluconazole (E). Each panel shows surface view of receptor-ligand complex, close view of receptor-ligand complex, Ribbon diagram of receptor showing green residues interacting with ligand (red), and LigPlot of receptor-ligand complex as indicated.

### 3.4 Drug likeness prediction of phytocompounds of *R. emodi*

The drug likeness filters helps in the early preclinical development by avoiding costly late step preclinical and clinical failure. The drug likeness properties of molecules were analyzed based on the Lipinski rule of 5. It was found that all the selected phytocompounds and antibiotics followed Lipinski’s rule of five except erythromycin) and amphotericin B (Table 5).

**Table 5:**
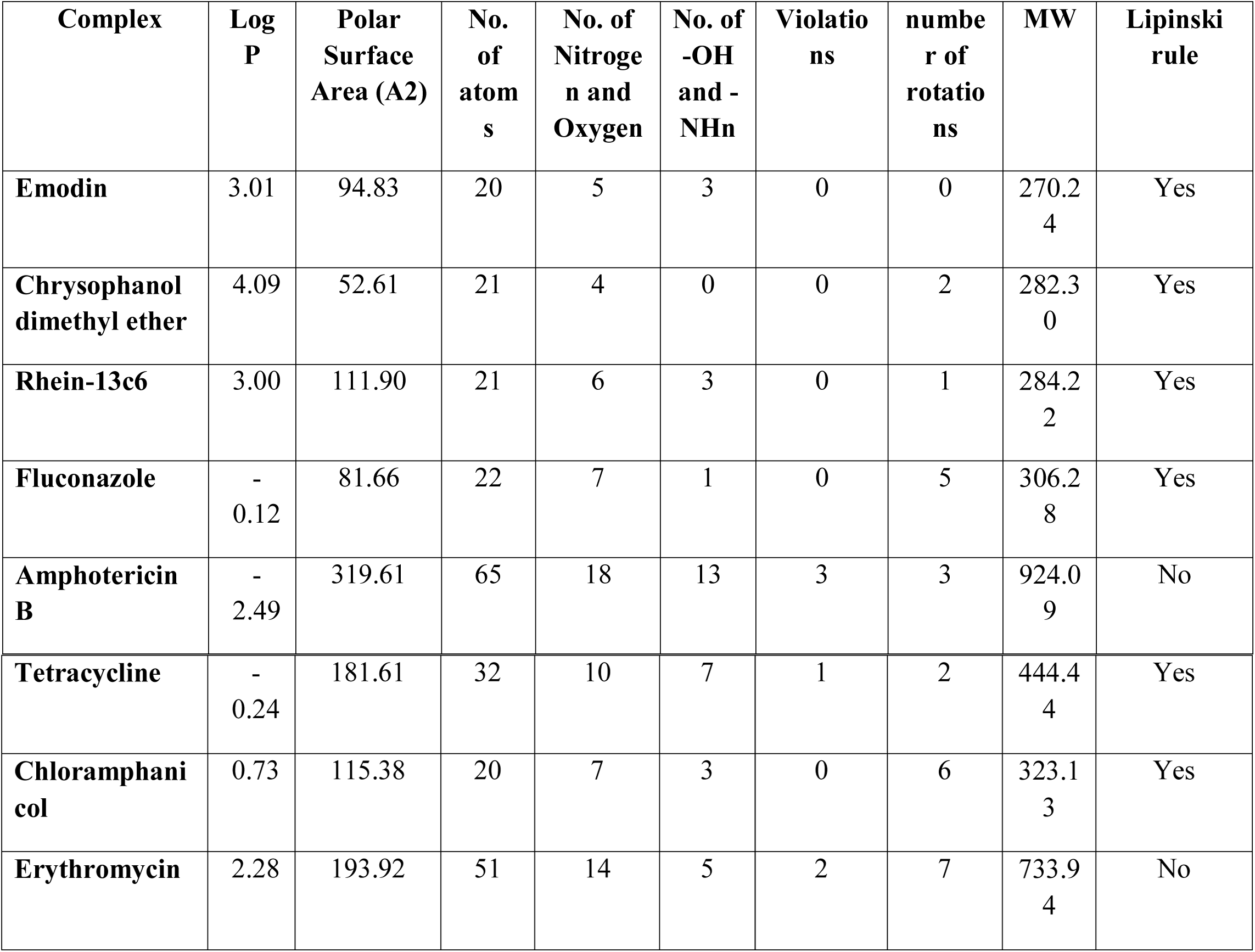
Drug-likeness prediction of selected phytocompounds from *R. emodi*.

| Complex | Log P | Polar Surface Area (A2) | No. of atoms | No. of Nitrogen and Oxygen | No. of -OH and -NHn | Violations | number of rotations | MW | Lipinski rule |
| --- | --- | --- | --- | --- | --- | --- | --- | --- | --- |
| **Emodin** | 3.01 | 94.83 | 20 | 5 | 3 | 0 | 0 | 270.24 | Yes |
| **Chrysophanol dimethyl ether** | 4.09 | 52.61 | 21 | 4 | 0 | 0 | 2 | 282.30 | Yes |
| **Rhein-13c6** | 3.00 | 111.90 | 21 | 6 | 3 | 0 | 1 | 284.22 | Yes |
| **Fluconazole** | -0.12 | 81.66 | 22 | 7 | 1 | 0 | 5 | 306.28 | Yes |
| **Amphotericin B** | -2.49 | 319.61 | 65 | 18 | 13 | 3 | 3 | 924.09 | No |
| **Tetracycline** | -<br>0.24 | 181.61 | 32 | 10 | 7 | 1 | 2 | 444.4<br>4 | Yes |
| **Chloramphenicol** | 0.73 | 115.38 | 20 | 7 | 3 | 0 | 6 | 323.1<br>3 | Yes |
| **Erythromycin** | 2.28 | 193.92 | 51 | 14 | 5 | 2 | 7 | 733.9<br>4 | No |

### 3.5 Toxicity and ADMET prediction of phytocompounds of *R. emodi*

Toxicity of phytocompounds was analyzed through Protox II server. The server admetSAR generates pharmacokinetic properties of compounds under different criteria: Absorption, Distribution, Metabolism, and Excretion (Cheng et al., 2012). The results of admetSAR analysis and toxicity prediction have been shown in table 5. All of the phytochemicals showed an acceptable range of ADMET profiles that reflect their efficiency as potent drug candidates. All the compounds showed good human intestinal solubility (HIA), except amphotericin B and erythromycin. Emodin showed similar acute rat toxicity (LD50) to that of antibacterial and antifungal drugs. None of the compounds are carcinogenic (Table-5). All the selected phytocompounds are inactive for cytotoxicity and hepatic toxicity. LD50 value for all selected compounds was higher, indicating non-toxic nature of these compounds. Among all these compounds, emodin was found to be safest as compared to that of both antibacterial and antifungal drugs. Thus, emodin fulfills all the enlisted criteria and we can suggest that it can be developed as potential antibacterial, and antifungal candidates for the development of a better drug.

**Table-5:** ADMET and Protox-II prediction of selected phytocompounds of *R. emodi* and drugs used through Admet SAR and Protox-II software.

| Compounds | Admet SAR |  |  | Protox-II |  |  |
| --- | --- | --- | --- | --- | --- | --- |
|  | Human intestinal absorption | Carcinogens | Rate Acute toxicity (LD50) kg/mol | LD<sub>50</sub>, (mg/kg) | Hepatotoxicity | Cytotoxicity |
| Emodin | + | Non-carcinogen | 2.5826 (III) | 5000 (class 5) | Inactive | Inactive |
| Rhein-13c6 | + | Non-carcinogen | 2.7118 (II) | 5000 (class 5) | Inactive | Inactive |
| Chrysophanol dimethyl ether | + | Non-carcinogen | 2.305 (II) | 5000 (class 5) | Inactive | Inactive |
| Fluconazole | + | Non-carcinogen | 2.407 (IV) | 1271 (Class 4) | Active | Inactive |
| Amphotericin B | - | Non-carcinogen | 2.577 (III) | 100 (class 3) | Inactive | Inactive |
| Chloramphenicol | + | Non-carcinogen | 1.676 (III) | 1500 (Class 4) | Inactive | Inactive |
| Erythromycin | - | Non-carcinogen | 3.136 (III) | 2000 (Class 4) | Active | Inactive |
| Tetracycline | + | Non-carcinogen | 3.011 (III) | 4400 (class4) | Active | Inactive |

## 4. Discussion

Computational strategies have gained an intense value in pharmaceutical research due to their ability to identify and develop novel promising compounds especially by molecular docking technique (Lounnas *et al*., 2013; Yuriev and Ramsland, 2013). Scientists from various research groups have applied these techniques to identify potential novel compounds against a variety of diseases (Ferreira *et al.*, 2015). In the present investigation, molecular docking studies was used to identify interactions between the major phytocompounds of *R. emodi* (Emodin, chrysophanol dimethyl ether and Rhein-13C6) (Rolta *et al.*, 2020) with antibacterial and antifungal receptor proteins. Standard antibacterial agents (chloramphanicol, tetracycline and erythromycin) and antifungal agents (fluconazole and amphotericin B) were used as control. Our study showed that Emodin, chrysophanol dimethyl ether and Rhein-13C6 are effective in term of their binding affinity or pharmacokinetic properties. Wuthi-Udomlert *et al.* (2010) reported the antifungal activity of anthraquinones (rhein and aloe emodin); while study from Rolta *et al.* (2020) reported the antibacterial, antifungal and synergistic activity of emodin isolated from chloroform sub-fraction of methanolic extract of *R. emodi* rhizome. Similar to our reports, Ahmed and Shohael (2019) also used *in silico* technique to identify the antifungal activity of aloe emodin, Chrysophanol, aloe-emodin and rhein from of *Senna alata* with Lanosterol 14 alpha demethylase (CYP51) target protein and they found aloe-emodin (−7.81 kcal mol^-1^), chrysophanol (−7.493 kcal mol^-1^) and rhein (−8.518 kcal mol^-1^) showed higher docking score than the native drug fluconazole (−6.856 kcalmol^-1^). Chrysophanol, aloe-emodin, rhein and emodin also fulfill the all criteria of Lipinski’s rule of five and ADMET. Tripathi *et al.* (2019) reported the emodin from *Aloe vera* can be used as potential therapeutics of cancer by molecular docking studies. Shadrack and Ndesendo (2017) evaluated emodin derivatives as inhibitors of Arylamine N-Acetyltransferase 2 (NAT2), Cyclooxygenase 2 and Topoisomerase 1 (TOP1) enzymes for colon and other forms of cancer. Docking studies suggested that D8 to be a target inhibitor of TOP1 while D5, D6 and D9 targets inhibitors of NAT2 enzymes. Pharmacokinetics suggested that these compounds can be potential anticancer agents. Physicochemical parameter correlated to the compounds activities. Sreelakshmi *et al.* (2017) have reported the strong binding affinities of kaempferol, chrysophanol and emodin identified from *Cassia tora* with epidermal growth factor receptor and validated the anti-cataractogenic potential of *C. tora* leaves.

## 5. Conclusion

To understand the mechanisms of synergistic potential of phytocompounds, the present study provides evidence that phytocompounds of *R. emodi* binds to bacterial and fungal proteins responsible for modifying the antibiotics and protecting the antibiotics and hence increase the potency. The *In silico* validation provide direct evidence to our *in vitro* study, where we reported that phytocompounds of *R. emodi* act synergistically in combination with antibacterial and antifungal antibiotics and lowered the dosage of antibiotics by 4–257 folds (Rolta *et al.*, 2020). Further, *In silico* study provide additional properties such as drug-likeness, ADMET prediction and toxicity analysis, which could help in developing non-toxic and effective combination formulation of phytocompounds and antibiotics.

## Acknowledgements

The authors acknowledge Shoolini University, Solan, for providing infrastructure support to conduct the research work. Authors also acknowledge the support provided by Yeast Biology Laboratory, School of Biotechnology, Shoolini University, Solan, India.

## Conflict of interest

Authors have no conflicts of interest

## Notes

### Competing Interest Statement

The authors have declared no competing interest.

